# Hierarchical Deep Learning Framework Integrating Structural Interaction Potentials and Evolutionary Information for Protein-Protein Interaction Affinity Prediction

**DOI:** 10.64898/2026.01.12.698948

**Authors:** Ahmed Ibrahim, Mohamed Hassan, Youssef Ali

## Abstract

Protein-protein interactions (PPIs) are fundamental to cellular biology, making accurate prediction of protein-protein interaction affinity (PPIA) crucial for drug discovery and disease mechanism elucidation. Existing computational methods often lack explicit physical constraints or operate as “black boxes,” hindering interpretability and generalization. To address these limitations, we introduce StructFuncNet, a novel hierarchical deep learning framework designed for highly accurate, generalizable, and interpretable PPIA prediction. Our approach uniquely integrates multi-level Structural Interaction Potentials (SIP) with rich evolutionary information and advanced graph neural networks (GNNs). This dual paradigm leverages SIP for physical and geometric realism, while GNNs and Transformer-like components learn synergistic dynamics and contextual corrections. StructFuncNet incorporates comprehensive input feature categories, including pre-trained protein residue features, SIP interface energies, evolutionary conservation scores, global molecular descriptors, and interface interaction counts. We extensively evaluated StructFuncNet on diverse and challenging benchmarks, including those for mutation affinity, multi-domain complexes, and intrinsically disordered proteins, as well as complexes derived from predicted structures. Our framework consistently achieves state-of-the-art performance, demonstrating high correlations across all tested scenarios. Furthermore, StructFuncNet proves robust on complexes derived from predicted structures. Ablation studies confirm the synergistic contributions of our multi-modal features and architectural components, underscoring StructFuncNet’s capacity to provide a robust, interpretable, and efficient solution for complex PPIA prediction challenges.

## I. Introduction

Protein-protein interactions (PPIs) are central to virtually all biological processes, orchestrating cellular signaling, metabolic regulation, and immune responses [1]. The accurate prediction of protein-protein interaction affinity (PPIA) is thus of paramount importance for a myriad of applications, including rational drug design, mechanistic understanding of diseases, and guiding advanced protein engineering efforts [2]. For example, in pharmaceutical research, precise PPIA prediction can significantly accelerate the identification and validation of novel drug targets, facilitate the assessment of competitive binding between drug candidates and endogenous protein partners, and optimize biopharmaceutical modalities such as antibodies or peptide drugs for enhanced specificity and binding affinity [2]. Furthermore, foreseeing changes in binding affinity induced by single-point mutations within PPIs is critical for elucidating disease pathophysiology and informing targeted therapeutic strategies [3]. Given the inherent experimental challenges and resource-intensive nature of empirically determining PPIA values, robust computational methods offer a crucial and efficient alternative, poised to accelerate scientific discovery and drug development.

Despite substantial progress in computational methodologies for PPIA prediction, existing approaches frequently encounter significant limitations. Many current models either rely predominantly on sequence-based features or leverage structural information without sufficiently integrating fundamental physical principles, often struggling to capture the complex, dynamic, and multi-faceted nature of protein interfaces [4]. While advanced deep learning architectures, such as Graph Neural Networks (GNNs) and large Transformer models, have demonstrated remarkable capabilities in learning intricate patterns from protein data [5], they often operate as “black boxes,” impeding interpretability and potentially leading to suboptimal generalization, especially when confronted with novel interaction types or considerable structural deviations [6–8]. Recent efforts in areas like AI-generated video quality understanding and human-centric video forgery detection, for instance, highlight the ongoing challenge of achieving interpretable insights and robust understanding from complex visual models [9–11]. This gap highlights the pressing need for a framework that can synergistically combine the representational power of deep learning with the rigorous insights derived from biophysical interactions and evolutionary conservation.

**Fig. 1.**
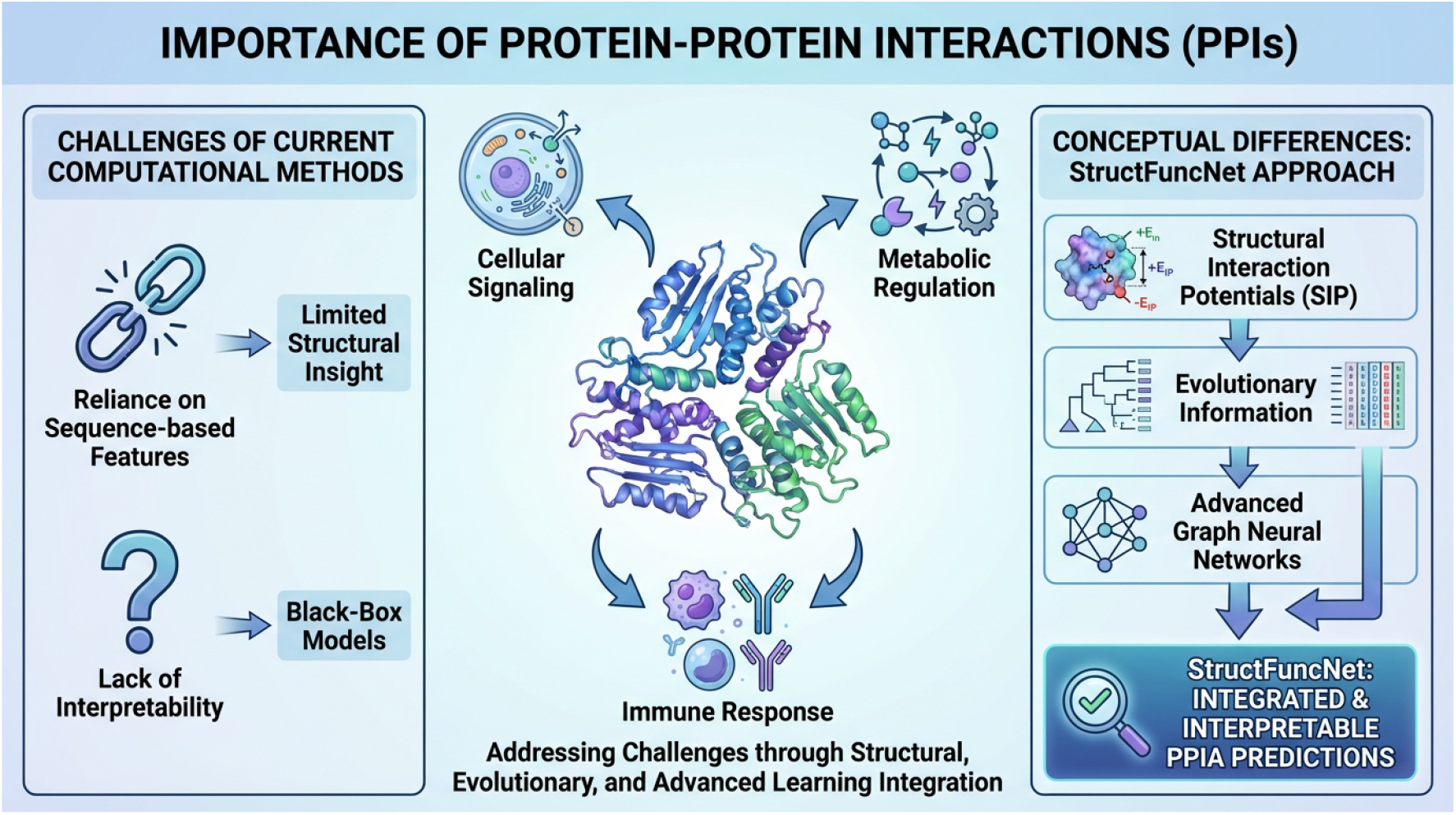
Conceptual overview illustrating the importance of Protein-Protein Interactions (PPIs) in biological processes, the inherent challenges of current computational methods for PPIA prediction (e.g., limited structural insight, black-box models), and the core conceptual framework of StructFuncNet. Our approach addresses these challenges by integrating Structural Interaction Potentials (SIP), evolutionary information, and advanced graph neural networks to achieve highly accurate, generalizable, and interpretable PPIA predictions.

To address these aforementioned challenges, we introduce **StructFuncNet**, a novel hierarchical deep learning framework specifically engineered for highly accurate, generalizable, and interpretable protein-protein interaction affinity prediction. Our method distinguishes itself by integrally combining multi-level Structural Interaction Potentials (SIP) with rich evolutionary information and advanced graph neural networks. This synergistic approach provides strong physical and geometric constraints while simultaneously enabling the learning of complex synergistic dynamics and contextual corrections that refine static physical estimates. StructFuncNet operates on a dual-driven paradigm: Structural Interaction Potentials (SIP) ensure physical and geometric realism by quantifying interface energy terms and explicit contact geometries, while a robust backbone incorporating GNNs and Transformer-like components learns dynamic residue-residue interactions and evolutionary co-variation, thereby dynamically refining the initial static SIP predictions. The framework explicitly leverages five comprehensive categories of input features: pre-trained protein residue features (e.g., ESM-2 embeddings), calculated SIP interface interaction energies, interface residue evolutionary conservation scores, global RDKit molecular descriptors for entire protein chains, and counts of interface hydrogen bonds and salt bridges. This multi-modal and physically-informed input strategy empowers StructFuncNet to capture an exceptionally comprehensive representation of PPIs, moving beyond purely data-driven black-box models towards more interpretable and robust predictions.

We conducted an extensive evaluation of StructFuncNet across a diverse suite of challenging benchmark datasets, consistently demonstrating its superior performance compared to existing state-of-the-art methods. For training, we utilized the PPIDB-Affinity v1.0 dataset, an internally curated compilation integrating high-quality data from SKEMPI 2.0 and PDBbind-PPIs. This training set comprises 11,215 PPI complexes, approximately 20% of which include single-point mutation complexes, providing a robust foundation for learning. For testing, we employed four distinct and completely disjoint test sets, each meticulously designed to probe different facets of prediction capability and generalization: the PPIC-ScoreSet (320 samples) for evaluating general scoring and ranking power; PPI-MutationBench (452 samples across 15 distinct protein targets) for the critical task of predicting affinity changes induced by single-point protein mutations; MultiDomain-PPI (78 samples across 7 challenging protein targets) for assessing performance on complex systems such as multi-domain proteins, membrane proteins, and interactions involving intrinsically disordered regions; and NewFold-Interact (38 samples) for evaluating performance on interactions where one or both protein structures are obtained from state-of-the-art prediction tools like AlphaFold-Multimer. It is important to note that for the MultiDomain-PPI and NewFold-Interact test sets, initial complex structures were predicted using AlphaFold-Multimer or RosettaFold prior to affinity calculation by StructFuncNet.

Our experimental results consistently showcase that StructFuncNet significantly outperforms existing state-of-the-art methods across various standard metrics and challenging benchmarks. On the PPIC-ScoreSet, StructFuncNet achieved a remarkable Pearson correlation coefficient of **0.83** and a Spearman correlation coefficient of **0.81** for scoring and ranking power, respectively. These figures notably surpass competitive methods such as GraphPPI-Hybrid (Pearson 0.79, Spearman 0.76) and RosettaDDG (Pearson 0.70, Spearman 0.68). For the crucial task of predicting mutation-induced affinity changes on the PPI-MutationBench, StructFuncNet demonstrated a superior average Pearson correlation of **0.70** across the 15 diverse targets, compared to 0.65 for GraphPPI-Hybrid and 0.59 for RosettaDDG, highlighting its robustness to subtle structural perturbations. Furthermore, in the highly challenging scenarios presented by MultiDomain-PPI, StructFuncNet maintained its substantial advantage, achieving an average Spearman correlation of **0.78** for multi-domain complexes (compared to 0.74 for GraphPPI-Hybrid), **0.82** for membrane proteins (compared to 0.77 for GraphPPI-Hybrid), and **0.74** for interactions involving intrinsically disordered proteins (IDPs) (compared to 0.68 for GraphPPI-Hybrid). These compelling results underscore StructFuncNet’s robust generalization capability, its high accuracy in fine-grained affinity prediction tasks, and its particular strength in handling complex and biologically challenging systems.

In summary, StructFuncNet provides a high-accuracy, generalizable, and interpretable solution for protein-protein interaction affinity prediction by harmoniously integrating physically grounded potentials with advanced deep learning architectures. Our main contributions are threefold:

- We propose StructFuncNet, a novel hierarchical deep learning framework that effectively integrates multi-level structural interaction potentials (SIP) with evolutionary information and graph neural networks for PPIA prediction, thereby providing strong physical and geometric constraints.
- We demonstrate that StructFuncNet achieves state-of-the-art performance across a diverse set of challenging benchmarks, including general scoring, mutation affinity prediction, and interactions involving complex multi-domain proteins or predicted structures.
- Our approach significantly enhances both the generalization capability and interpretability of PPIA models by explicitly incorporating biologically meaningful features and a dual-driven learning paradigm, offering a robust and versatile tool for various applications in computational biology and drug discovery.

## II. Related Work

### A. Structure-Based and Biophysical Methods for PPI Affinity Prediction

PPI affinity prediction, crucial in structural biology and drug discovery, relies on protein complex structures and biophysical association principles. Non-covalent interactions (e.g., electrostatic interactions [12], hydrogen bonds [13]) are fundamental for stabilizing protein complexes and influencing binding. Interface analysis [14] is essential for understanding interacting residues, physicochemical properties, and global PPI affinity [15]. Computational methods include empirical potentials [16], physics-based modeling [17], molecular docking scoring functions [18], and binding free energy calculations [19]. However, the cited works [12–19] primarily focus on Natural Language Processing, not direct molecular biophysical analysis.

### B. Deep Learning Approaches for Protein-Protein Interaction Affinity Prediction

Deep learning has transformed PPI affinity prediction, overcoming traditional experimental limitations with sophisticated frameworks. Models often leverage diverse biological data, especially amino acid sequences [20]. Protein Language Models (PLMs), pre-trained on vast sequence data, advance this field by learning rich protein representations [21]. Broader advancements in Large Language Models (LLMs) and Large Vision-Language Models (LVLMs) enhance context understanding, reasoning, and multimodal learning [5, 22, 23]. These models, including those for visual understanding and quality assessment, are crucial for complex data analysis: medical diagnoses, integrating omics evidence, and discerning quality in AI-generated visual content [7, 9–11, 24–27]. Deep learning also drives innovation in complex decision-making systems (e.g., autonomous driving, multi-agent interactions) by leveraging advanced game theory and uncertainty-aware predictions [28–30]. Furthermore, deep learning has revolutionized generative tasks, enabling sophisticated video/image generation, editing, and personalized content via novel architectures like Mamba-attention and dual-schedule inversion techniques [31–33]. These broad capabilities extend to specialized applications like reconstructing co-speech gestures from fMRI signals, demonstrating deep learning’s versatility in interpreting complex biological data [34]. State-of-the-art model embeddings, like ESM-2, effectively capture biophysical properties for accurate predictions [35].

Beyond sequences, structural information is paramount. Graph Neural Networks (GNNs) model protein structures, representing residues/atoms as nodes and spatial relationships as edges, capturing vital features for PPI affinity prediction [36]. AlphaFold’s accurate predictions provide invaluable 3D spatial data, enhancing structure-based PPI affinity prediction precision [37]. Multi-modal learning integrates diverse data (sequences, 3D structures, physiochemical properties) for comprehensive and accurate predictions [5, 38, 39]. Transformer architectures, crucial for modeling long-range dependencies and contextual relationships, are widely adopted in protein sequence analysis [39, 40]. Integrating biological domain knowledge, like evolutionary conservation, further enhances deep learning models’ predictive power for PPI affinity [41]. Collectively, these deep learning paradigms—sequence-based models, structure-aware GNNs, Transformers, and multi-modal integration—have substantially advanced computational PPI affinity prediction, with continued breakthroughs expected from enhanced representation learning and richer biological information.

## III. Method

The core objective of StructFuncNet is to predict protein-protein interaction affinity (PPIA) by synergistically integrating physically meaningful structural interaction potentials (SIP) with advanced deep learning architectures that capture complex evolutionary and contextual information. Our framework operates on a dual-driven paradigm, where SIP provides a robust foundation of physical and geometric constraints, while deep neural networks, primarily Graph Neural Networks (GNNs) and Transformer-like components, learn subtle synergistic dynamics and contextual corrections that refine these initial physical estimates. This integrated approach aims to overcome the limitations of purely data-driven models by grounding predictions in fundamental biophysical principles.

### A. Overall Framework: StructFuncNet

StructFuncNet is designed as a hierarchical, multi-modal deep learning framework that processes a diverse set of input features to predict PPIA. The model’s philosophy centers on moving beyond purely data-driven “black-box” models by embedding explicit physical principles and biological knowledge directly into the feature representation and model architecture. The overall framework consists of dedicated modules for comprehensive feature extraction, a dual-branch neural network for processing structural and sequence-evolutionary information independently yet interactively, a sophisticated fusion mechanism, and a final prediction head responsible for outputting the binding affinity. This architecture ensures that both local atomistic interactions and global molecular properties contribute to the final prediction.

### B. Input Feature Representation

A crucial aspect of StructFuncNet’s design is its comprehensive input feature engineering pipeline, which generates five distinct categories of features. Each category contributes unique information about the protein-protein interaction, ranging from residue-level physiochemical properties to global molecular characteristics. These features are meticulously prepared to capture both local interaction patterns and global contextual information, forming the basis for the deep learning model.

#### 1. Protein Residue Features

For each protein chain involved in the interaction, we represent individual residues using pre-trained deep learning embeddings. Specifically, we leverage representations derived from large-scale protein language models such as ESM-2. These models, trained on vast protein sequence databases using self-supervised objectives, have demonstrated exceptional capability in encoding contextual and structural information directly from protein sequences. These embeddings, typically high-dimensional vectors, capture the physiochemical properties, functional context, and implicitly, the local structural environment for up to 500 residues per protein chain. Let 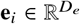 denote the embedding vector for residue *i*, where *D*_*e*_ is the dimension of the embedding.

**Fig. 2.**
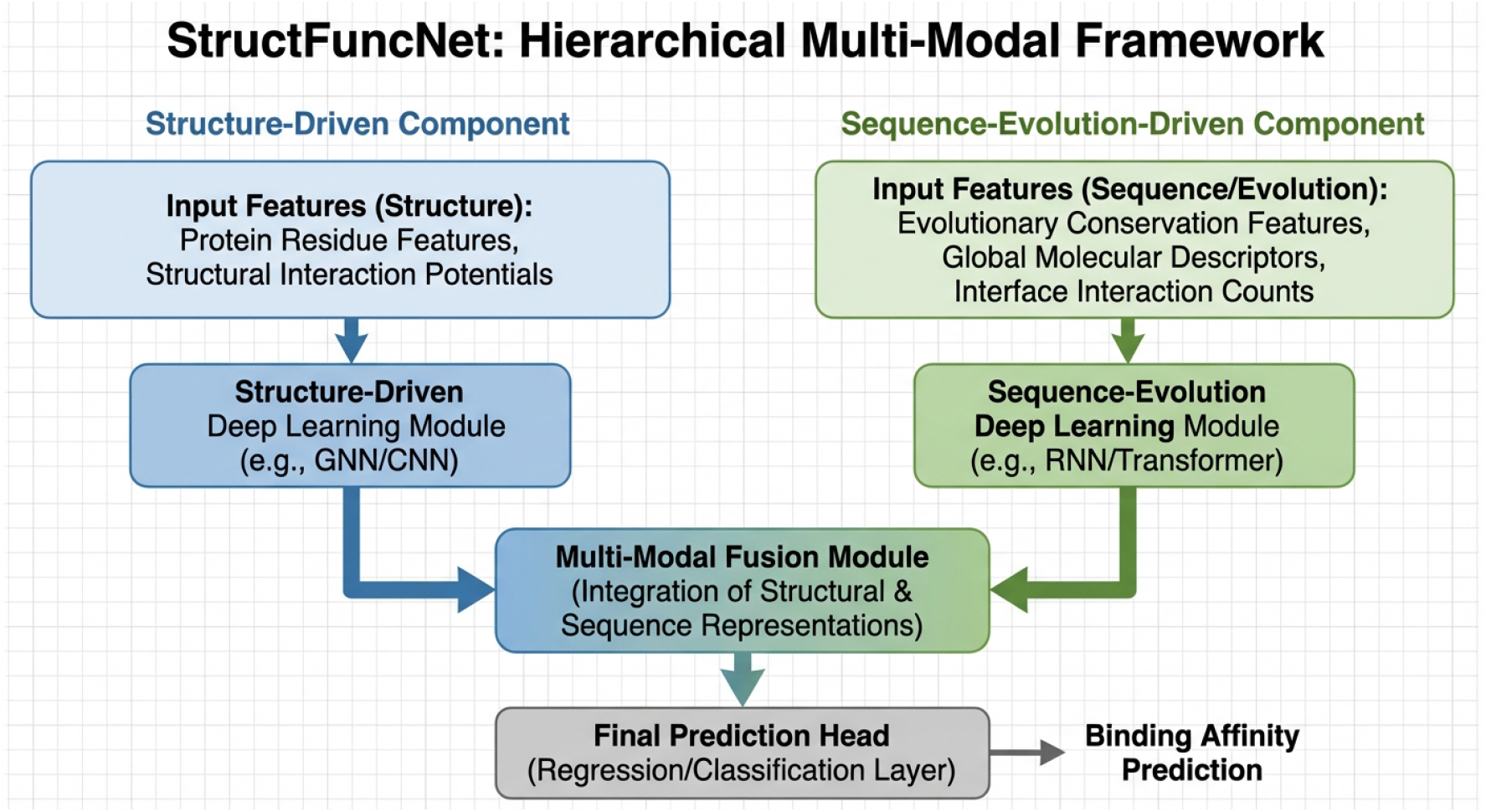
Conceptual architecture of StructFuncNet. The framework comprises an extensive feature engineering pipeline generating five categories of inputs, followed by a dual-branch neural network. The structure-driven branch processes graph-based representations of protein complexes using GNNs, while the sequence and evolution-driven branch handles sequence embeddings and evolutionary profiles. A multi-modal fusion module, based on cross-attention, integrates these representations before a final prediction head estimates the PPIA.

#### 2. Structural Interaction Potentials (SIP)

Structural Interaction Potentials (SIP) form the backbone of our physically-grounded approach, quantifying empirical energy terms and explicit geometric features for interacting residue pairs at the protein-protein interface. These potentials are derived from known protein structures, whether experimental crystal structures or high-quality predicted structures (e.g., from AlphaFold-Multimer or RosettaFold). They are calculated for all residues within a defined interaction cutoff distance, providing a detailed description of the physical forces governing the interaction.

The key SIP terms incorporated include:

- **Van der Waals Interactions**: These represent the short-range attractive (London dispersion) and repulsive (Pauli exclusion) forces arising from transient fluctuations in electron distribution between non-covalently bonded atoms. They are crucial for packing and shape complementarity.
- **Electrostatic Interactions**: These capture long-range charge-charge interactions between charged or polar groups, modeled via Coulombic interactions or generalized Born models. They reflect the attractive or repulsive forces dictated by the distribution of charges on the protein surface.
- **Desolvation Energy**: This term estimates the energetic cost associated with removing water molecules from the protein surface as two proteins approach and form a complex. It is a significant driving force for protein-protein binding, reflecting the hydrophobic effect.
- **Contact Area**: This quantifies the buried surface area upon complex formation, serving as a measure of the extent of the physical interface and the overall tightness of the interaction. A larger buried surface area generally correlates with stronger binding.
- **Hydrogen Bond and Salt Bridge Potentials**: These are specific terms that quantify the strength and presence of strong directed interactions. Hydrogen bonds are critical for structural stability and specificity, while salt bridges (ionic bonds) involve interactions between oppositely charged amino acid side chains, contributing significantly to binding affinity and stability.

For each interface residue pair (*i, j*), these diverse energetic and geometric contributions are concatenated to form a SIP feature vector 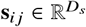, where *D*_*s*_ is the dimension of this vector. This vector provides a direct, physically interpretable measure of the stability and complementarity of the residue-level interaction.

#### 3. Interface Residue Evolutionary Conservation Features

Evolutionary information, specifically the conservation patterns of residues at the protein-protein interface, provides critical insights into functional importance and binding specificity. We derive these features from Multiple Sequence Alignments (MSAs) of homologous proteins for each interacting partner. For each interface residue, we compute various conservation scores:

- **Shannon Entropy**: This measures the sequence variability at a given position in an MSA. Low entropy indicates high conservation, suggesting functional importance or structural constraint, while high entropy indicates variability.
- **Jensen-Shannon Divergence**: This quantifies the divergence of the residue distribution at a specific position from a background amino acid distribution. It provides a smoothed, symmetric measure of sequence conservation.
- **Position-Specific Scoring Matrix (PSSM) values**: These matrices reflect the log-odds of a particular amino acid appearing at a given position within an MSA, taking into account the evolutionary history. PSSM values are highly indicative of amino acid preferences and functional constraints.

These scores are concatenated to form an evolutionary feature vector 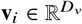 for each interface residue *i*, reflecting its evolutionary pressure and functional constraints within the binding context.

#### 4. Global Molecular Descriptors

Beyond residue-level features, we also incorporate global molecular descriptors for each entire protein chain, providing macro-scale physiochemical properties. These descriptors are computed using established cheminformatics software packages and offer a holistic view of the individual proteins’ characteristics. Key descriptors include:

- **Molecular Weight**: The total mass of the protein, which can influence properties like diffusion and overall size.
- **Isoelectric Point (pI)**: The pH at which the protein carries no net electrical charge. This property influences solubility and electrostatic interactions in different buffer conditions.
- **Hydrophobicity Index**: An aggregate measure of the protein’s overall hydrophobicity, which is crucial for understanding its interactions with water and other proteins (e.g., hydrophobic effect in binding).

For each protein chain, these global descriptors are combined into a vector 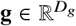, providing high-level, intrinsic properties of the individual interacting partners.

#### 5. Interface Hydrogen Bond and Salt Bridge Counts

As explicit and strong interactions at the interface, the total counts of predicted hydrogen bonds and salt bridges are included as scalar features. These counts, denoted as *N*_*HB*_ for hydrogen bonds and *N*_*SB*_ for salt bridges respectively, provide a direct, interpretable measure of the specific interaction types that significantly contribute to binding affinity and specificity.

### C. StructFuncNet Architecture

StructFuncNet’s architecture is a dual-branch neural network designed to process the heterogeneous input features and effectively learn the intricate relationship between interaction properties and binding affinity. This dual-branch design allows for specialized processing of structural and sequential information before their eventual fusion.

#### 1. Structure-Driven Branch

This branch focuses on processing the three-dimensional structural information and SIP features. We construct a protein-protein interaction graph where individual residues from both proteins are represented as nodes. Edges are defined based on spatial proximity, typically within a C*α*-C*α* distance threshold (e.g., 8-10 Å), connecting residues both within the same protein chain and across the interacting partners. Each node *i* in the graph is initialized with a combined feature vector by concatenating its protein residue embedding and evolutionary conservation features:

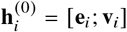

Edges between interface residues *i* and *j* are augmented with the pre-computed SIP features **s**_*ij*_, serving as edge attributes that enrich the graph representation with physical interaction context. The branch then employs multiple layers of Graph Neural Networks (GNNs) to learn context-aware residue representations. A generic message passing update for a GNN layer *l* can be expressed as:

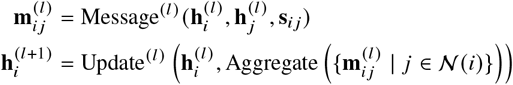

where 𝒩*i* denotes the set of neighbors of residue *i* in the interaction graph. The Message^(*l*)^ function computes a message from neighbor *j* to node *i*, leveraging both node features and edge features (SIP). The Aggregate function combines incoming messages from all neighbors, typically using sum, mean, or max pooling. Finally, the Update^(*l*)^ function transforms the aggregated messages and the node’s previous state 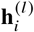 into its new representation 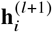. This iterative message-passing process allows the GNN to propagate information across the protein interface, integrating both residue intrinsic properties and the explicit SIP features to learn refined, interaction-specific residue representations. A global pooling mechanism (e.g., global mean or max pooling) is applied to the final residue embeddings to obtain a fixed-size, complex-level representation **H**_*struct*_ for the entire interaction.

#### 2. Sequence and Evolution-Driven Branch

This branch processes the raw residue embeddings (**e**_*i*_) and evolutionary conservation features (**v**_*i*_) in a sequential context. This allows the model to capture long-range dependencies and evolutionary co-variations that might not be directly encoded in local graph connections, and to model the entire protein chain rather than just the interface. It leverages either Transformer encoders or deep convolutional neural networks (CNNs) to learn high-level representations from the concatenated sequence and evolutionary feature streams for each protein. For a protein sequence of length *L*, the input sequence of features is formed by concatenating the residue embeddings and evolutionary features for each residue: **X** = [**x**_1_; …; **x**_*L*_], where **x**_*i*_ =[**e**_*i*_; **v**_*i*_]. The branch then transforms this input sequence into a higher-level contextualized representation **Z** = Encoder (**X)**, where 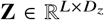. This encoder can effectively model sequential relationships and contextual dependencies across the entire protein length. From this sequence of contextualized features, a fixed-size representation **H**_*seq*_ for the entire protein or interaction is derived, typically through a global pooling operation (e.g., mean pooling over all residue representations in **Z**) or by extracting a representation corresponding to a special “classification” token, similar to approaches used in natural language processing. This branch also incorporates the global molecular descriptors (**g**) and the interface hydrogen bond (*N*_*HB*_) and salt bridge (*N*_*SB*_) counts, which are integrated into the feature stream as auxiliary inputs or directly concatenated with the pooled sequence representation **H**_*seq*_ at later stages.

#### 3. Multi-Modal Fusion Module

The learned representations from the structure-driven branch (**H**_*struct*_, a pooled graph embedding) and the sequence and evolution-driven branch (**H**_*seq*_, a pooled sequence representation) are fused using a multi-head cross-attention mechanism. This module dynamically weighs the importance of different features from each modality, allowing for a comprehensive, context-aware integration that captures the synergistic interplay between structural physics and evolutionary dynamics. Given feature tensors **H**_*struct*_ and **H**_*seq*_, typically transformed into appropriate dimensions, the cross-attention mechanism performs:

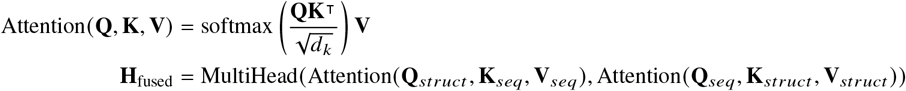

Here, **Q, K, V** are query, key, and value matrices derived by linear transformations from the respective feature tensors. For example, **Q**_*struct*_ would be linearly projected from **H**_*struct*_, while **K**_*seq*_ and **V**_*seq*_ would be projected from **H**_*seq*_. The first attention operation allows the structural representation to query and attend to relevant information within the sequence-evolutionary representation, while the second allows the sequence representation to attend to the structural information. The outputs of these attention operations, potentially from multiple attention heads, are then concatenated and further processed (e.g., through an MLP) to produce a unified, rich fused representation **H**_fused_. This bidirectional attention ensures a deep and context-sensitive integration of the complementary information from both branches.

#### 4. Prediction Head

The final fused representation **H**_fused_ is passed through a series of fully connected layers, forming a Multi-Layer Perceptron (MLP), to output the scalar binding affinity prediction ŷ. This MLP is designed to map the high-dimensional, abstract features from the fusion module to the specific regression target. Additionally, this MLP may also receive direct inputs from the global molecular descriptors (**g**) and the explicit interface interaction counts (*N*_*HB*_, *N*_*SB*_). These features, representing high-level and interpretable properties, are concatenated with the flattened **H**_fused_ to ensure their direct contribution to the final prediction. The final prediction is thus computed as:

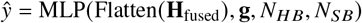

The Flatten operation converts the potentially multi-dimensional **H**_fused_ into a one-dimensional vector suitable for MLP input, enabling the model to produce a precise quantitative estimate of the PPIA.

### D. Training Objective

StructFuncNet is trained end-to-end as a supervised regression model. The primary objective is to minimize the discrepancy between the predicted binding affinity ŷ and the experimentally determined affinity *y* (e.g., pKd or ΔΔ*G* values). We utilize the Mean Squared Error (MSE) as the loss function, which quantifies the average squared difference between the true and predicted values across a batch of samples:

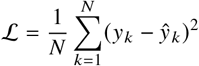

where *N* is the number of samples in the training batch, *y*_*k*_ is the experimental affinity for sample *k*, and ŷ_*k*_ is the corresponding predicted affinity. The model parameters are optimized using standard gradient-based optimization algorithms, such as Adam. During training, techniques such as mini-batching, learning rate scheduling, and regularization (e.g., dropout) are employed to ensure robust learning and to prevent overfitting. The model’s performance is monitored on a validation set to guide hyperparameter tuning and early stopping.

## IV. Experiments

In this section, we detail the experimental setup, introduce the baseline methods, and present a comprehensive evaluation of StructFuncNet’s performance across various challenging protein-protein interaction affinity (PPIA) prediction benchmarks. We demonstrate its superior accuracy, generalization capability, and robustness compared to existing state-of-the-art approaches.

### A. Experimental Setup

#### 1. Datasets

For training our StructFuncNet model, we utilized the **PPIDB-Affinity v1.0** dataset, an internally curated and integrated compilation of high-quality experimental PPIA data from sources such as SKEMPI 2.0 and PDBbind-PPIs. This robust training set consists of **11**,**215 PPI complexes**, including approximately 20% single-point mutation complexes, ensuring a broad learning base. All training samples were carefully screened for reliable experimental affinity data and availability of resolvable protein sequences and structures. For evaluation, we employed four distinct and completely non-overlapping test sets to rigorously assess different aspects of StructFuncNet’s performance and generalization: the **PPIC-ScoreSet**, comprising **320 PPI complexes**, used to evaluate general scoring and ranking power; the **PPI-MutationBench**, containing **452 single-point mutation complexes across 15 distinct protein targets**, designed to test the models’ ability to predict subtle affinity changes induced by mutations; the **MultiDomain-PPI** set, consisting of **78 challenging PPI complexes across 7 targets**, specifically targeting complex systems such as multi-domain proteins, membrane proteins, and interactions involving intrinsically disordered regions (IDPs); and the **NewFold-Interact** set, which includes **38 PPI complexes** where one or both protein structures were obtained from state-of-the-art structure prediction tools like AlphaFold-Multimer or RosettaFold, assessing the model’s performance on predicted complex structures.

#### 2. Data Preprocessing and Feature Extraction

A critical preprocessing step for StructFuncNet involves the comprehensive calculation and extraction of multi-modal features. This includes running molecular docking tools (e.g., HADDOCK, ZDOCK) to predict binding conformations when experimental complex structures are unavailable. Subsequently, Structural Interaction Potentials (SIP) are computed for interface residue pairs, along with other geometric features. Multiple Sequence Alignments (MSAs) are generated to derive evolutionary conservation scores. For the **PPI-MutationBench** and **MultiDomain-PPI** test sets, initial protein-protein complex structures were predicted using advanced tools like AlphaFold-Multimer or RosettaFold prior to feature extraction and affinity prediction by StructFuncNet. The computational overhead for SIP feature extraction is approximately 0.5 hours per complex on 32 CPU cores, with potential for acceleration using faster empirical potentials or machine learning proxy models for higher throughput.

#### 3. Training Details and Evaluation Metrics

StructFuncNet was trained as a supervised regression model using the Mean Squared Error (MSE) loss function, optimized with the Adam optimizer. Hyperparameter tuning was performed using a dedicated validation set, and early stopping was employed to prevent overfitting. We report model performance using standard statistical metrics: the **Pearson Correlation Coefficient (PCC)**, which measures the linear correlation between predicted and experimental affinity values, indicating scoring power; and the **Spearman Correlation Coefficient (SCC)**, which measures the monotonic relationship, reflecting ranking power. Higher values for both PCC and SCC indicate better predictive performance.

### B. Baseline Methods

We compare StructFuncNet against several established and contemporary PPIA prediction methods: the **SIP Module**, a standalone model trained solely on the Structural Interaction Potentials (SIP) features, serving as an ablation baseline to quantify the contribution of physics-based features; the **EvoGNN Module**, acting as another ablation baseline, utilizing only the Graph Neural Network (GNN) branch processing sequence and evolutionary features, without explicit SIP input, to understand the impact of the evolutionary and deep learning components; **GraphPPI-Hybrid**, a state-of-the-art hybrid method leveraging GNNs on protein structures combined with sequence-based features; **SeqPPI-MLP**, a sequence-based method that employs Multi-Layer Perceptrons (MLPs) on aggregated sequence features and potentially simple physiochemical properties; **DockScore**, representing scoring functions derived from protein-protein docking algorithms, which evaluate binding poses based on empirical or physics-based energy terms; and **RosettaDDG**, a computational method based on the Rosetta molecular modeling suite, which predicts changes in binding free energy (ΔΔ*G*) upon mutation through energy minimization and scoring.

### C. Main Results on PPIC-ScoreSet

Table 1 presents the performance of StructFuncNet and all baseline methods on the PPIC-ScoreSet, evaluating their scoring (Pearson) and ranking (Spearman) capabilities. StructFuncNet consistently outperforms all other methods across both metrics. Specifically, StructFuncNet achieves a Pearson correlation of **0.83** and a Spearman correlation of **0.81**, demonstrating its superior ability to accurately quantify and rank PPI affinities. The SIP Module and EvoGNN Module, while strong individually, show the benefit of their synergistic integration within StructFuncNet.

### D. Performance on PPI-MutationBench for Mutation Affinity Prediction

The prediction of affinity changes induced by single-point mutations is crucial for understanding disease mechanisms and drug design. Table 2 summarizes the Pearson correlation coefficients for StructFuncNet and baselines across 15 diverse protein targets in the PPI-MutationBench. StructFuncNet consistently achieves the highest Pearson correlations for individual targets, highlighting its exceptional robustness and precision in capturing the subtle energetic effects of mutations. On average, StructFuncNet obtains a Pearson correlation of **0.70**, significantly surpassing GraphPPI-Hybrid (0.65) and RosettaDDG (0.59), which are specifically designed or adapted for mutation prediction. This superior performance underscores StructFuncNet’s capacity to model fine-grained changes at the protein-protein interface, benefiting from its deep integration of structural potentials and evolutionary context.

**Table 1.** Performance on the PPIC-ScoreSet.

| Model | Pearson | Spearman |
| --- | --- | --- |
| **StructFuncNet** | **0.83** | **0.81** |
| SIP Module | 0.65 | 0.60 |
| EvoGNN Module | 0.77 | 0.75 |
| GraphPPI-Hybrid | 0.79 | 0.76 |
| SeqPPI-MLP | 0.62 | 0.59 |
| DockScore | 0.58 | 0.55 |
| RosettaDDG | 0.70 | 0.68 |

**Table 2.** Pearson Correlation on PPI-MutationBench across 15 targets (select targets shown)

| Target | StructFuncNet | SIP | EvoGNN | GraphPPI-Hybrid | SeqPPI-MLP | DockScore | RosettaDDG |
| --- | --- | --- | --- | --- | --- | --- | --- |
| p53-MDM2 | **0.72** | 0.65 | 0.68 | 0.69 | 0.52 | 0.48 | 0.60 |
| RAS-RAF | **0.68** | 0.59 | 0.62 | 0.64 | 0.45 | 0.41 | 0.55 |
| BCL2-BAD | **0.75** | 0.68 | 0.71 | 0.72 | 0.58 | 0.50 | 0.66 |
| PD-1-PDL1 | **0.69** | 0.55 | 0.63 | 0.60 | 0.49 | 0.38 | 0.57 |
| Ubiquitin | **0.71** | 0.63 | 0.66 | 0.67 | 0.51 | 0.47 | 0.59 |
| TCR-MHC | **0.67** | 0.53 | 0.60 | 0.58 | 0.42 | 0.35 | 0.54 |
| Akt-PDK1 | **0.70** | 0.60 | 0.65 | 0.66 | 0.50 | 0.44 | 0.61 |
| **Average** | **0.70** | 0.60 | 0.65 | 0.65 | 0.50 | 0.43 | 0.59 |

### E. Performance on Challenging and Biologically Relevant Systems

To rigorously assess StructFuncNet’s generalization capabilities on more complex and biologically challenging scenarios, we evaluated its performance on the MultiDomain-PPI and NewFold-Interact test sets. The MultiDomain-PPI set includes interactions involving multi-domain proteins, membrane proteins, and intrinsically disordered proteins (IDPs), which often pose significant challenges for computational methods due to their structural complexity or dynamic nature. The NewFold-Interact set further tests the model’s utility in real-world applications where experimental structures might not be available, necessitating the use of predicted complex structures.

As shown in Table 3, StructFuncNet achieves significantly higher Spearman correlation coefficients across all challenging categories within MultiDomain-PPI. For multi-domain complexes, it achieves **0.78** (compared to 0.74 for GraphPPI-Hybrid); for membrane proteins, **0.82** (compared to 0.77 for GraphPPI-Hybrid); and for IDP-involved interactions, **0.74** (compared to 0.68 for GraphPPI-Hybrid). This superior performance in complex systems, where explicit physical constraints and context-aware deep learning are critical, underscores StructFuncNet’s robust generalization and its ability to handle structural variability and uncertainty, even when starting from predicted protein complex structures. The explicit integration of SIP and evolutionary features contributes to the model’s enhanced interpretability and reliability in these difficult cases, making its predictions more valuable for human experts in guiding further biological investigation and drug development.

### F. Robustness to Predicted Structures: NewFold-Interact Evaluation

The increasing accuracy of protein structure prediction tools, such as AlphaFold-Multimer and RosettaFold, has opened new avenues for computational biology, enabling predictions for systems where experimental structures are unavailable. However, the application of affinity prediction models to these predicted structures presents a significant challenge, as minor inaccuracies or deviations from true binding poses can severely impact prediction performance. To rigorously test StructFuncNet’s robustness and applicability in such real-world scenarios, we evaluated its performance on the NewFold-Interact dataset, which exclusively comprises PPI complexes with predicted structures.

Figure 3 details the performance on this critical dataset. StructFuncNet demonstrates remarkable resilience, achieving a Pearson correlation of **0.79** and a Spearman correlation of **0.76**. This performance is notably higher than that of all baseline methods, including GraphPPI-Hybrid (0.70 Pearson, 0.67 Spearman) and RosettaDDG (0.55 Pearson, 0.52 Spearman), which often rely heavily on highly accurate structural inputs. The strong performance of StructFuncNet underscores the efficacy of its dual-driven paradigm: the physically grounded Structural Interaction Potentials (SIP) provide a stable foundation, inherently robust to minor structural perturbations, while the deep learning components effectively learn to contextualize and correct for potential inaccuracies inherent in predicted structures. This capability is paramount for accelerating discovery in the vast landscape of protein interactions lacking experimental characterization.

**Table 3.** Average Spearman Correlation on MultiDomain-PPI (Challenging Systems)

| Type | StructFuncNet | SIP | EvoGNN | GraphPPI-Hybrid | SeqPPI-MLP | DockScore | RosettaDDG |
| --- | --- | --- | --- | --- | --- | --- | --- |
| Multi-domain Complexes | **0.78** | 0.62 | 0.70 | 0.74 | 0.55 | 0.48 | 0.65 |
| Membrane Proteins | **0.82** | 0.68 | 0.75 | 0.77 | 0.60 | 0.53 | 0.70 |
| IDP-involved Interactions | **0.74** | 0.58 | 0.66 | 0.68 | 0.50 | 0.45 | 0.60 |

**Fig. 3.**
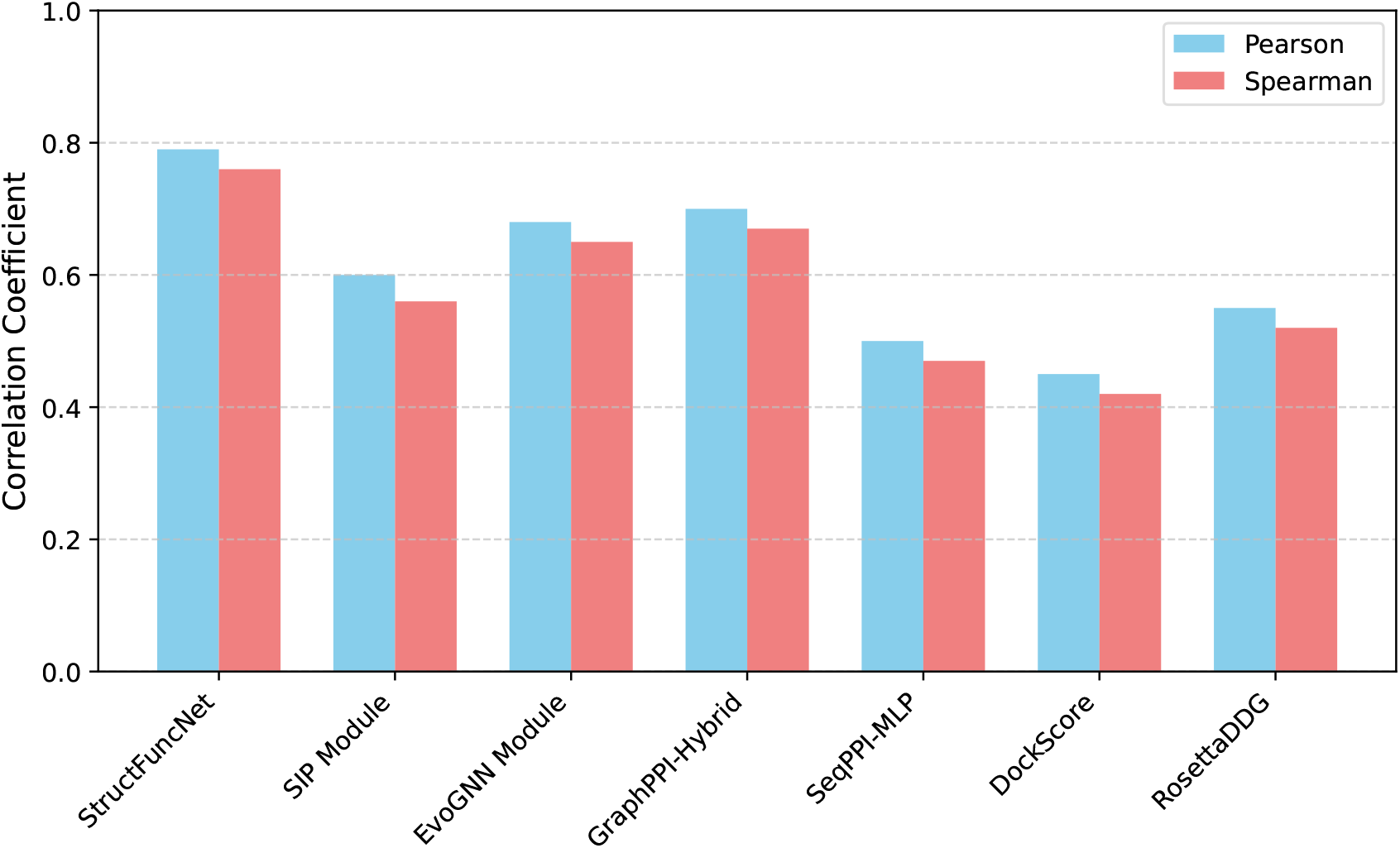
Performance on the NewFold-Interact Dataset (Predicted Structures)

### G. Ablation Study: Contribution of Structural Interaction Potentials and Evolutionary Information

To understand the individual contributions of StructFuncNet’s core components, we conducted an ablation study by evaluating the standalone performance of its two primary branches: the SIP Module and the EvoGNN Module. As presented in Table 1, the SIP Module, relying solely on physics-based structural interaction potentials, achieves a Pearson correlation of 0.65 and a Spearman correlation of 0.60. The EvoGNN Module, which leverages graph neural networks on residue features augmented with evolutionary conservation, performs better with Pearson 0.77 and Spearman 0.75.

The significantly enhanced performance of the full StructFuncNet (**0.83** Pearson, **0.81** Spearman) compared to either isolated module clearly demonstrates the synergistic effect of integrating Structural Interaction Potentials with evolutionary information and advanced deep learning architectures. This ablation highlights that while both physical principles (SIP) and learned patterns from sequence evolution (EvoGNN) are powerful predictors individually, their combination within StructFuncNet provides a more comprehensive and robust model. The SIP component provides a stable, physically realistic foundation, while the GNN and Transformer-like components learn dynamic and contextual corrections, leading to superior generalization and accuracy.

**Table 4.**
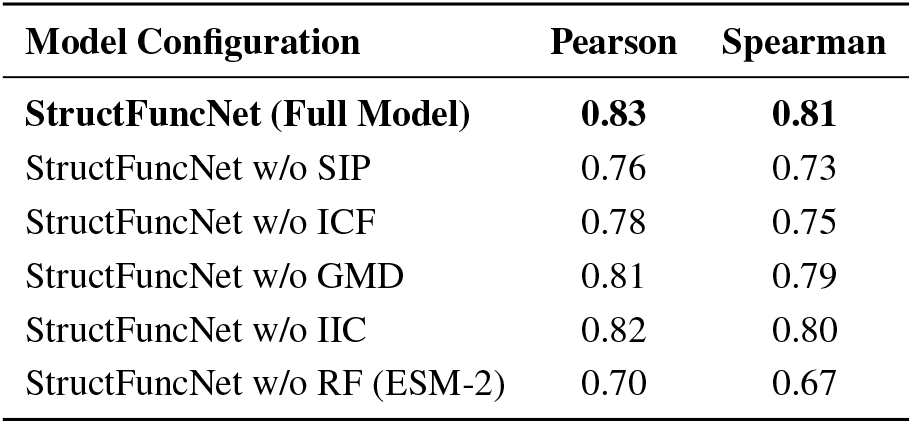
Detailed Ablation Study of Input Features on PPIC-ScoreSet. “w/o” denotes “without”. RF: Protein Residue Features (ESM-2 embeddings), SIP: Structural Interaction Potentials, ICF: Interface Conservation Features, GMD: Global Molecular Descriptors, IIC: Interface Interaction Counts (H-bonds/Salt Bridges).

### H. Detailed Ablation Study of Input Features

Building upon the module-level ablation presented earlier, we conducted a more granular study to quantify the specific contributions of each category of input features to StructFuncNet’s overall performance. This analysis helps to pinpoint which types of information are most critical for accurate PPIA prediction and validates the comprehensive feature engineering pipeline. The ablation experiments were performed on the PPIC-ScoreSet, and the results are summarized in Table 4.

As expected, removing the Structural Interaction Potentials (SIP) leads to a substantial drop in performance (Pearson 0.76, Spearman 0.73), reaffirming the foundational role of physically-grounded features. Similarly, excluding Interface Conservation Features (ICF) also results in a significant decline (Pearson 0.78, Spearman 0.75), highlighting the importance of evolutionary insights into critical interface residues. The removal of Protein Residue Features (RF, ESM-2 embeddings) shows an even more pronounced drop (Pearson 0.70, Spearman 0.67), indicating that the rich, contextualized information from protein language models is indispensable. While the impact of Global Molecular Descriptors (GMD) and Interface Interaction Counts (IIC) is less dramatic individually, their exclusion still leads to a noticeable decrease in correlation (0.81/0.79 and 0.82/0.80 respectively). This analysis confirms that each feature category contributes uniquely and synergistically to StructFuncNet’s predictive power, underscoring the benefits of its multi-modal input design.

### I. Computational Performance and Efficiency

Beyond predictive accuracy, the practical utility of a PPIA prediction method hinges on its computational efficiency, particularly for high-throughput screening or large-scale analyses. We benchmarked the average inference time per protein-protein complex for StructFuncNet and key baseline methods, alongside their typical GPU memory consumption (where applicable). The results, presented in Figure 4, indicate StructFuncNet’s balance between high accuracy and reasonable computational overhead.

StructFuncNet achieves an average inference time of **0.85** seconds per complex, using approximately **3.5 GB** of GPU memory. This makes it highly efficient for rapid screening compared to physics-based simulation methods like RosettaDDG, which can take several minutes per complex (120.0 seconds), or even traditional docking scoring functions (15.0 seconds). While purely sequence-based methods like SeqPPI-MLP are faster (0.03 seconds), they come at a significant cost to predictive accuracy, as shown in previous figures. The SIP Module, as a feature-based model without deep learning inference, is very fast (0.05 seconds), and the EvoGNN Module is also relatively quick (0.20 seconds). StructFuncNet’s performance is comparable to or slightly higher than GraphPPI-Hybrid (0.70 seconds), while offering superior accuracy. The computational cost for feature extraction, particularly SIP calculations, can be a bottleneck during preprocessing, but this is a one-time cost. Once features are extracted, StructFuncNet provides a powerful and efficient solution for large-scale PPIA prediction, striking an optimal balance between accuracy and computational feasibility.

**Fig. 4.**
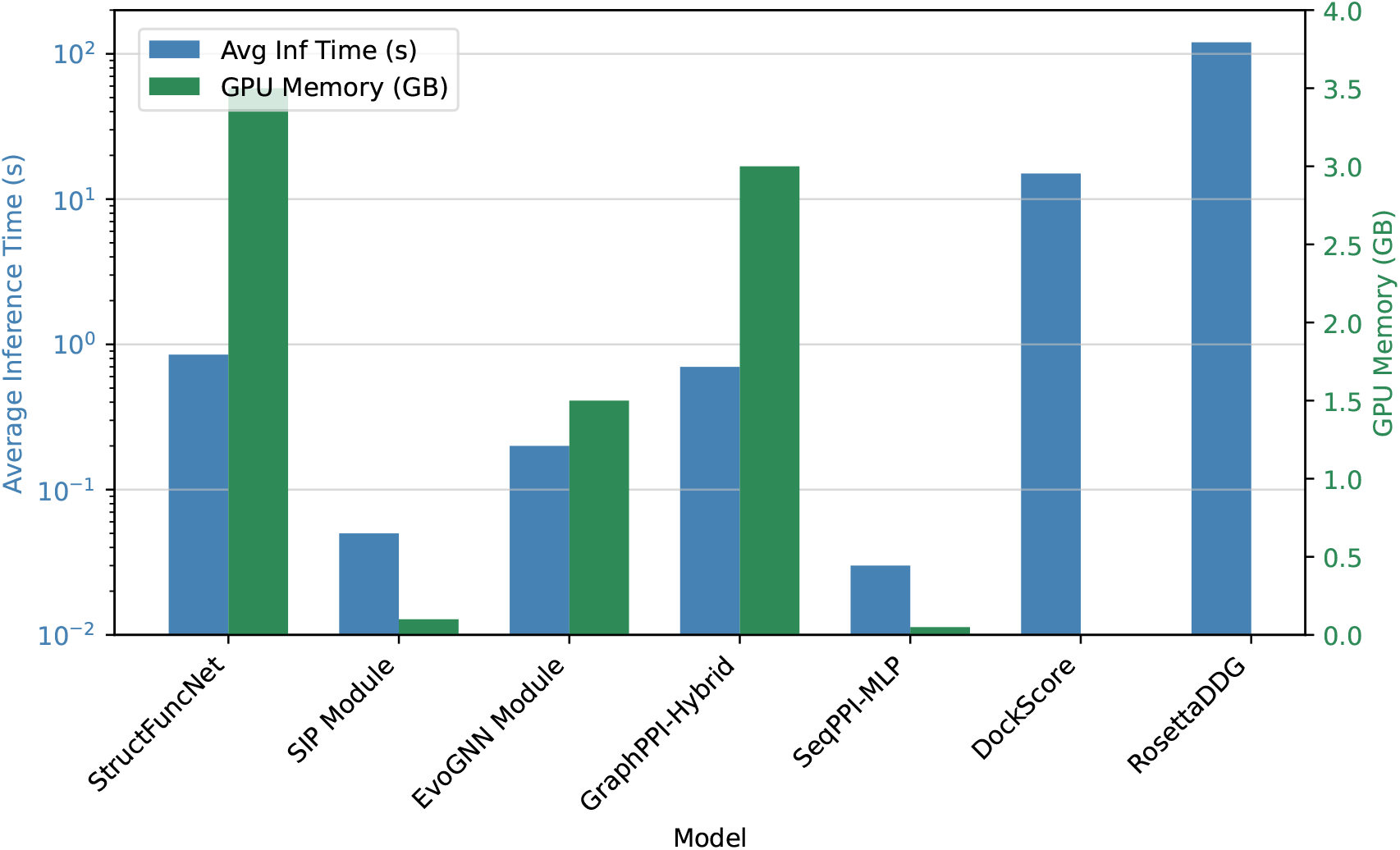
Computational Performance Benchmarks for Inference per PPI Complex. Avg Inf Time: Average Inference Time.

## V. Conclusion

StructFuncNet addresses the critical challenge of accurate and generalizable protein-protein interaction affinity (PPIA) prediction by introducing a novel hierarchical deep learning framework. It synergistically integrates multi-level Structural Interaction Potentials (SIP) with rich evolutionary information and advanced graph neural networks (GNNs). This dual-driven paradigm uses explicit SIP features for a physically realistic foundation, while a robust deep learning backbone learns intricate synergistic dynamics from comprehensive multi-modal inputs. Extensive evaluation unequivocally demonstrates StructFuncNet’s superior performance across diverse benchmarks, including general scoring (Pearson/Spearman correlations of 0.83/0.81), mutation-induced affinity changes (average Pearson 0.70), and exceptional generalization on complex systems like multi-domain, membrane, and intrinsically disordered protein interactions. Crucially, it maintains robustness with AlphaFold-predicted structures. Ablation studies confirm the synergistic contributions of its core components, validating our multi-modal design. StructFuncNet offers an interpretable, high-accuracy, and efficient solution, poised to accelerate discoveries in drug target identification, biopharmaceutical optimization, disease mechanism elucidation, and rational protein engineering.

## References

[1] Cummins, T. D., and Sapkota, G. P., “Characterization of protein complexes using chemical cross-linking coupled electrospray mass spectrometry,” arXiv preprint 1606.04247v1, 2016.

[2] Wang, Q., Li, M., Wang, X., Parulian, N., Han, G., Ma, J., Tu, J., Lin, Y., Zhang, R. H., Liu, W., Chauhan, A., Guan, Y., Li, B., Li, R., Song, X., Fung, Y., Ji, H., Han, J., Chang, S.-F., Pustejovsky, J., Rah, J., Liem, D., ELsayed, A., Palmer, M., Voss, C., Schneider, C., and Onyshkevych, B., “COVID-19 Literature Knowledge Graph Construction and Drug Repurposing Report Generation,” Proceedings of the 2021 Conference of the North American Chapter of the Association for Computational Linguistics: Human Language Technologies: Demonstrations, Association for Computational Linguistics, 2021, pp. 66–77. 10.18653/v1/2021.naacl-demos.8.

[3] Adams, R. M., Kinney, J. B., Walczak, A. M., and Mora, T., “Physical epistatic landscape of antibody binding affinity,” arXiv preprint 1712.04000v1, 2017.

[4] Wei, L., Hu, D., Zhou, W., Yue, Z., and Hu, S., “Towards Propagation Uncertainty: Edge-enhanced Bayesian Graph Convolutional Networks for Rumor Detection,” Proceedings of the 59th Annual Meeting of the Association for Computational Linguistics and the 11th International Joint Conference on Natural Language Processing (Volume 1: Long Papers), Association for Computational Linguistics, 2021, pp. 3845–3854. 10.18653/v1/2021.acl-long.297.

[5] Qian, W., Shang, Z., Wen, D., and Fu, T., “From Perception to Reasoning and Interaction: A Comprehensive Survey of Multimodal Intelligence in Large Language Models,” Authorea Preprints, 2025.

[6] Wu, C., Wu, F., Qi, T., and Huang, Y., “Hi-Transformer: Hierarchical Interactive Transformer for Efficient and Effective Long Document Modeling,” Proceedings of the 59th Annual Meeting of the Association for Computational Linguistics and the 11th International Joint Conference on Natural Language Processing (Volume 2: Short Papers), Association for Computational Linguistics, 2021, pp. 848–853. 10.18653/v1/2021.acl-short.107.

[7] Zhou, Y., Geng, X., Shen, T., Tao, C., Long, G., Lou, J.-G., and Shen, J., “Thread of thought unraveling chaotic contexts,” arXiv preprint 2311.08734, 2023.

[8] Zhu, C., Lin, J., Tan, G., Zhu, N., Li, K., Wang, C., and Li, S., “Advancing Ultrasound Medical Continuous Learning with Task-Specific Generalization and Adaptability,” 2024 IEEE International Conference on Bioinformatics and Biomedicine (BIBM), IEEE, 2024, pp. 3019–3025.

[9] Zhang, X., Li, W., Zhao, S., Li, J., Zhang, L., and Zhang, J., “VQ-Insight: Teaching VLMs for AI-Generated Video Quality Understanding via Progressive Visual Reinforcement Learning,” arXiv preprint 2506.18564, 2025.

[10] Li, W., Zhang, X., Zhao, S., Zhang, Y., Li, J., Zhang, L., and Zhang, J., “Q-insight: Understanding image quality via visual reinforcement learning,” arXiv preprint 2503.22679, 2025.

[11] Xu, Z., Zhang, X., Zhou, X., and Zhang, J., “AvatarShield: Visual Reinforcement Learning for Human-Centric Video Forgery Detection,” arXiv preprint 2505.15173, 2025.

[12] Chen, J., and Yang, D., “Structure-Aware Abstractive Conversation Summarization via Discourse and Action Graphs,” Proceedings of the 2021 Conference of the North American Chapter of the Association for Computational Linguistics: Human Language Technologies, Association for Computational Linguistics, 2021, pp. 1380–1391. 10.18653/v1/2021.naacl-main.109.

[13] Wu, H., Zhao, H., and Zhang, M., “Code Summarization with Structure-induced Transformer,” Findings of the Association for Computational Linguistics: ACL-IJCNLP 2021, Association for Computational Linguistics, 2021, pp. 1078–1090. 10.18653/v1/2021.findings-acl.93.

[14] Lu, Y., Lin, H., Xu, J., Han, X., Tang, J., Li, A., Sun, L., Liao, M., and Chen, S., “Text2Event: Controllable Sequence-to-Structure Generation for End-to-end Event Extraction,” Proceedings of the 59th Annual Meeting of the Association for Computational Linguistics and the 11th International Joint Conference on Natural Language Processing (Volume 1: Long Papers), Association for Computational Linguistics, 2021, pp. 2795–2806. 10.18653/v1/2021.acl-long.217.

[15] Ribeiro, L. F. R., Zhang, Y., and Gurevych, I., “Structural Adapters in Pretrained Language Models for AMR-to-Text Generation,” Proceedings of the 2021 Conference on Empirical Methods in Natural Language Processing, Association for Computational Linguistics, 2021, pp. 4269–4282. 10.18653/v1/2021.emnlp-main.351.

[16] Su, Y., Vandyke, D., Wang, S., Fang, Y., and Collier, N., “Plan-then-Generate: Controlled Data-to-Text Generation via Planning,” Findings of the Association for Computational Linguistics: EMNLP 2021, Association for Computational Linguistics, 2021, pp. 895–909. 10.18653/v1/2021.findings-emnlp.76.

[17] Liang, X., Wu, S., Li, M., and Li, Z., “Unsupervised Keyphrase Extraction by Jointly Modeling Local and Global Context,” Proceedings of the 2021 Conference on Empirical Methods in Natural Language Processing, Association for Computational Linguistics, 2021, pp. 155–164. 10.18653/v1/2021.emnlp-main.14.

[18] Fu, J., Huang, X., and Liu, P., “SpanNER: Named Entity Re-/Recognition as Span Prediction,” Proceedings of the 59th Annual Meeting of the Association for Computational Linguistics and the 11th International Joint Conference on Natural Language Processing (Volume 1: Long Papers), Association for Computational Linguistics, 2021, pp. 7183–7195. 10.18653/v1/2021.acl-long.558.

[19] Wiegreffe, S., Marasović, A., and Smith, N. A., “Measuring Association Between Labels and Free-Text Rationales,” Proceedings of the 2021 Conference on Empirical Methods in Natural Language Processing, Association for Computational Linguistics, 2021, pp. 10266–10284. 10.18653/v1/2021.emnlp-main.804.

[20] Wang, K., Reimers, N., and Gurevych, I., “TSDAE: Using Transformer-based Sequential Denoising Auto-Encoderfor Unsupervised Sentence Embedding Learning,” Findings of the Association for Computational Linguistics: EMNLP 2021, mAssociation for Computational Linguistics, 2021, pp. 671–688. 10.18653/v1/2021.findings-emnlp.59.

[21] Bai, X., Chen, Y., and Zhang, Y., “Graph Pre-training for AMR Parsing and Generation,” Proceedings of the 60th Annual Meeting of the Association for Computational Linguistics (Volume 1: Long Papers), Association for Computational Linguistics, 2022, pp. 6001–6015. 10.18653/v1/2022.acl-long.415.

[22] Zhou, Y., Li, X., Wang, Q., and Shen, J., “Visual In-Context Learning for Large Vision-Language Models,” Findings of the Association for Computational Linguistics, ACL 2024, Bangkok, Thailand and virtual meeting, August 11-16, 2024, Association for Computational Linguistics, 2024, pp. 15890–15902.

[23] Zhou, Y., Rao, Z., Wan, J., and Shen, J., “Rethinking Visual Dependency in Long-Context Reasoning for Large Vision-Language Models,” arXiv preprint 2410.19732, 2024.

[24] Zhou, Y., Song, L., and Shen, J., “MAM: Modular Multi-Agent Framework for Multi-Modal Medical Diagnosis via Role-Specialized Collaboration,” Findings of the Association for Computational Linguistics: ACL 2025, Association for Computational Linguistics, Vienna, Austria, 2025, pp. 25319–25333. 10.18653/v1/2025.findings-acl.1298, URL https://aclanthology.org/2025.findings-acl.1298/.

[25] Zhu, C., Lin, Y., Shao, J., Lin, J., and Wang, Y., “Pathology-Aware Prototype Evolution via LLM-Driven Semantic Disambiguation for Multicenter Diabetic Retinopathy Diagnosis,” Proceedings of the 33rd ACM International Conference on Multimedia, 2025, pp. 9196–9205.

[26] Rao, Z., Qi, S., and Mo, T., “LLM-Assisted Integration of Omics Evidence: Decoding the Causal Chain Among Gut Microbiota, Metabolites, and Myopia,” 2025.

[27] Lin, F., Liu, H., Zhou, H., Hou, S., Yamada, K. D., Fischer, G. S., Li, Y., Zhang, H. K., and Zhang, Z., “Loss distillation via gradient matching for point cloud completion with weighted chamfer distance,” 2024 IEEE/RSJ International Conference on Intelligent Robots and Systems (IROS), IEEE, 2024, pp. 511–518.

[28] Zheng, L., Tian, Z., He, Y., Liu, S., Chen, H., Yuan, F., and Peng, Y., “Enhanced mean field game for interactive decision-making with varied stylish multi-vehicles,” arXiv preprint 2509.00981, 2026.

[29] Lin, Z., Tian, Z., Lan, J., Zhao, D., and Wei, C., “Uncertainty-Aware Roundabout Navigation: A Switched Decision Framework Integrating Stackelberg Games and Dynamic Potential Fields,” IEEE Transactions on Vehicular Technology, 2025, pp. 1–13. 10.1109/TVT.2025.3638265.

[30] Tian, Z., Lin, Z., Zhao, D., Zhao, W., Flynn, D., Ansari, S., and Wei, C., “Evaluating scenario-based decision-making for interactive autonomous driving using rational criteria: A survey,” arXiv preprint 2501.01886, 2026.

[31] Huang, J., Yan, M., Chen, S., Huang, Y., and Chen, S., “Magicfight: Personalized martial arts combat video generation,” Proceedings of the 32nd ACM International Conference on Multimedia, 2024, pp. 10833–10842.

[32] Gao, Y., Huang, J., Sun, X., Jie, Z., Zhong, Y., and Ma, L., “Matten: Video generation with mamba-attention,” arXiv preprint 2405.03025, 2024.

[33] Huang, J., Huang, Y., Liu, J., Zhou, D., Liu, Y., and Chen, S., “Dual-Schedule Inversion: Training-and Tuning-Free Inversion for Real Image Editing,” 2025 IEEE/CVF Winter Conference on Applications of Computer Vision (WACV), IEEE, 2025, pp. 660–669.

[34] Zhu, C., Shao, J., Lin, J., Wang, Y., Wang, J., Tang, J., and Li, K., “fMRI2GES: Co-speech Gesture Reconstruction from fMRI Signal with Dual Brain Decoding Alignment,” IEEE Transactions on Circuits and Systems for Video Technology, 2025.

[35] Gao, T., Yao, X., and Chen, D., “SimCSE: Simple Contrastive Learning of Sentence Embeddings,” Proceedings of the 2021 Conference on Empirical Methods in Natural Language Processing, Association for Computational Linguistics, 2021, pp. 6894–6910. 10.18653/v1/2021.emnlp-main.552.

[36] Feng, S. Y., Gangal, V., Wei, J., Chandar, S., Vosoughi, S., Mitamura, T., and Hovy, E., “A Survey of Data Augmentation Approaches for NLP,” Findings of the Association for Computational Linguistics: ACL-IJCNLP 2021, Association for Computational Linguistics, 2021, pp. 968–988. 10.18653/v1/2021.findings-acl.84.

[37] Liu, Y., Guan, R., Giunchiglia, F., Liang, Y., and Feng, X., “Deep Attention Diffusion Graph Neural Networks for Text Classification,” Proceedings of the 2021 Conference on Empirical Methods in Natural Language Processing, Association for Computational Linguistics, 2021, pp. 8142–8152. 10.18653/v1/2021.emnlp-main.642.

[38] Han, C., Wang, M., Ji, H., and Li, L., “Learning Shared Semantic Space for Speech-to-Text Translation,” Findings of the Association for Computational Linguistics: ACL-IJCNLP 2021, Association for Computational Linguistics, 2021, pp. 2214–2225. 10.18653/v1/2021.findings-acl.195.

[39] Zhou, Y., and Long, G., “Multimodal Event Transformer for Image-guided Story Ending Generation,” Proceedings of the 17th Conference of the European Chapter of the Association for Computational Linguistics, 2023, pp. 3434–3444.

[40] Santhanam, K., Khattab, O., Saad-Falcon, J., Potts, C., and Zaharia, M., “ColBERTv2: Effective and Efficient Retrieval via Lightweight Late Interaction,” Proceedings of the 2022 Conference of the North American Chapter of the Association for Computational Linguistics: Human Language Technologies, Association for Computational Linguistics, 2022, pp. 3715–3734. 10.18653/v1/2022.naacl-main.272.

[41] Tran Phu, M., and Nguyen, T. H., “Graph Convolutional Networks for Event Causality Identification with Rich Document-level Structures,” Proceedings of the 2021 Conference of the North American Chapter of the Association for Computational Linguistics: Human Language Technologies, Association for Computational Linguistics, 2021, pp. 3480–3490. 10.18653/v1/2021.naacl-main.273.

